# Community standards to facilitate development and address challenges in metabolic modeling

**DOI:** 10.1101/700112

**Authors:** Maureen A. Carey, Andreas Dräger, Jason A. Papin, James T. Yurkovich

## Abstract

Standardization of data and models facilitates effective communication, especially in computational systems biology. However, both the development and consistent use of standards and resources remains challenging. As a result, the amount, quality, and format of the information contained within systems biology models are not consistent and therefore present challenges for widespread use and communication. Here, we focused on these standards, resources, and challenges in the field of metabolic modeling by conducting a community-wide survey. We used this feedback to (1) outline the major challenges that our field faces and to propose solutions and (2) identify a set of features that defines what a “gold standard” metabolic network reconstruction looks like concerning content, annotation, and simulation capabilities. We anticipate that this community-driven outline will help the long-term development of community-inspired resources as well as produce high-quality, accessible models. More broadly, we hope that these efforts can serve as blueprints for other computational modeling communities to ensure continued development of both practical, usable standards and reproducible, knowledge-rich models.

## INTRODUCTION

Systems biology uses holistic approaches to understand the networks that comprise biological systems. Computational models that attempt to represent these systems are inherently complex with many interacting components, requiring the mathematical formalization of biological phenomenon. Standardizing how these phenomena are represented is thus required to organize these complexities and make these formalizations interpretable and accessible. Many resources—including databases, algorithms, file formats, software, and compiled ‘best practices’—exist to facilitate standardization (Dräger and Palsson 2014; Le Novère *et al*. 2005; Waltemath *et al*. 2011; Hucka *et al*. 2003; Brazma *et al*. 2006; Ravikrishnan and Raman 2015; Stanford *et al*. 2015), but the consistent use and application of these standards can pose a significant challenge (Ebrahim *et al*. 2015).

Here, we use the term ‘standard’ to represent a framework or resource to facilitate interpretation and evaluate quality, particularly involving data formatting and presentation in computational modeling. Beyond merely evaluating quality (Mbacham *et al*. 2014), standardization can also improve efficiency (Rose *et al*. 2010), content (Waltemath *et al*. 2016), reproducibility (Ebrahim *et al*. 2015), code and model sharing (Yurkovich *et al*. 2017), and the ease of entering a field (Thiele and Palsson 2010). Much of the domain-specific knowledge embedded into a computational model can be uninterpretable to scientists other than the developer or inaccessible to those outside the field if unstandardized. This inefficiency limits useful discussion between biologists, modelers, software developers, and database architects and emphasizes the importance of researchers at the interface of these fields. Standardization, however, combats this inefficiency and makes a computational model interpretable and accessible.

The modeling process has two phases: model construction and simulation. Decisions about technical approaches and biological content to include in the model are made throughout both the construction and simulation processes, influencing the downstream use of the model. These implicit and explicit decisions affect the reusability; if the design decisions made during the model building process do not match well with a particular application, the quality of the simulation results will suffer. Such design decisions are influenced by a scientist’s perspective, a motivating biological question, and data availability, as well as a scientist’s familiarity with and access to existing resources. Manual steps of this process are particularly vulnerable to potential biases and thus are inherently irreproducible, emphasizing the role of diligent tracking of references and design decisions. Field-defined best practices and standards can help control for or evaluate quality.

Here, we discuss existing standards in computational biological modeling and when and why they are not met, building on previous efforts to assess standardization in computational systems biology (Stanford *et al*. 2015). While standards are not unique to any individual field, we use metabolic network modeling as a case study in which to discuss the challenges to accept and implement standards. We first discuss how metabolic models are built, reviewing existing standards and their application to metabolic modeling. Next, we highlight challenges that the metabolic modeling community faces in effectively utilizing these resources, identified from a community survey. Finally, we propose an integrated set of standards which we hope will serve as a checklist to ensure standardized content across reconstructions to improve accessibility, interpretability, and consistency. These standards are aimed at improving the quality of metabolic network reconstructions and lowering the activation energy required for biologists to build new reconstructions or use existing reconstructions. We hope that our proposed checklists will help increase the rigor of the field and the accessibility of metabolic modeling—for both experts and newcomers alike—as well as provide a model for sustainable standardization for other systems biology fields.

## STANDARDIZATION IN METABOLIC MODELING: A CASE STUDY

The metabolic modeling community frequently utilizes COnstraint-Based Reconstruction and Analysis (COBRA) methods to build and compute computational models that represent an organism’s metabolic phenotype. Investigating metabolic processes using mechanistic computational models has led to insights into many complex phenotypes and phenomena, including drug resistance (Dunphy and Papin 2018), aging (Nilsson *et al*. 2017), and cancer (Bordbar *et al*. 2014). The construction of genome-scale metabolic network reconstructions and models is a multi-step process that involves the reconstruction of a metabolic network, manual curation to incorporate known physiology, computation of metabolic phenotypes, and the distribution of the models and results (Box 1). The COBRA field has been led by community-driven, open-source software efforts (Ebrahim *et al*. 2013; Heirendt *et al*. 2019) developed to enable these kinds of analyses, building on existing systems biological standards and principles.

### Model structure

SBML—the *de facto* standard file format for sharing systems biology models (including metabolic models)—provides infrastructure for storing and sharing biological data and models (Hucka *et al*. 2003), and leading developers of SBML were integral to the development of MIRIAM and MIASE. SBML files encode biological models in a machine-readable format, support all MIRIAM-recommended data, and are the most common format for editing and sharing metabolic reconstructions (Fig 1). SBML files contain lists of system components with corresponding parameters linking these components (e.g., metabolites in a reaction) and constraints (e.g., compartmentalization, reaction bounds). Saving a reconstruction as an SBML file thus inherently reinforces a set of standards. Further, the SBML field also offers several model validators and a test suite to identify non-standard formatting in COBRA models (Table 1). Other formats for sharing models, such as SBtab (Lubitz *et al*. 2016), have also been developed for the sharing of systems biology models.

**Fig. 1.**
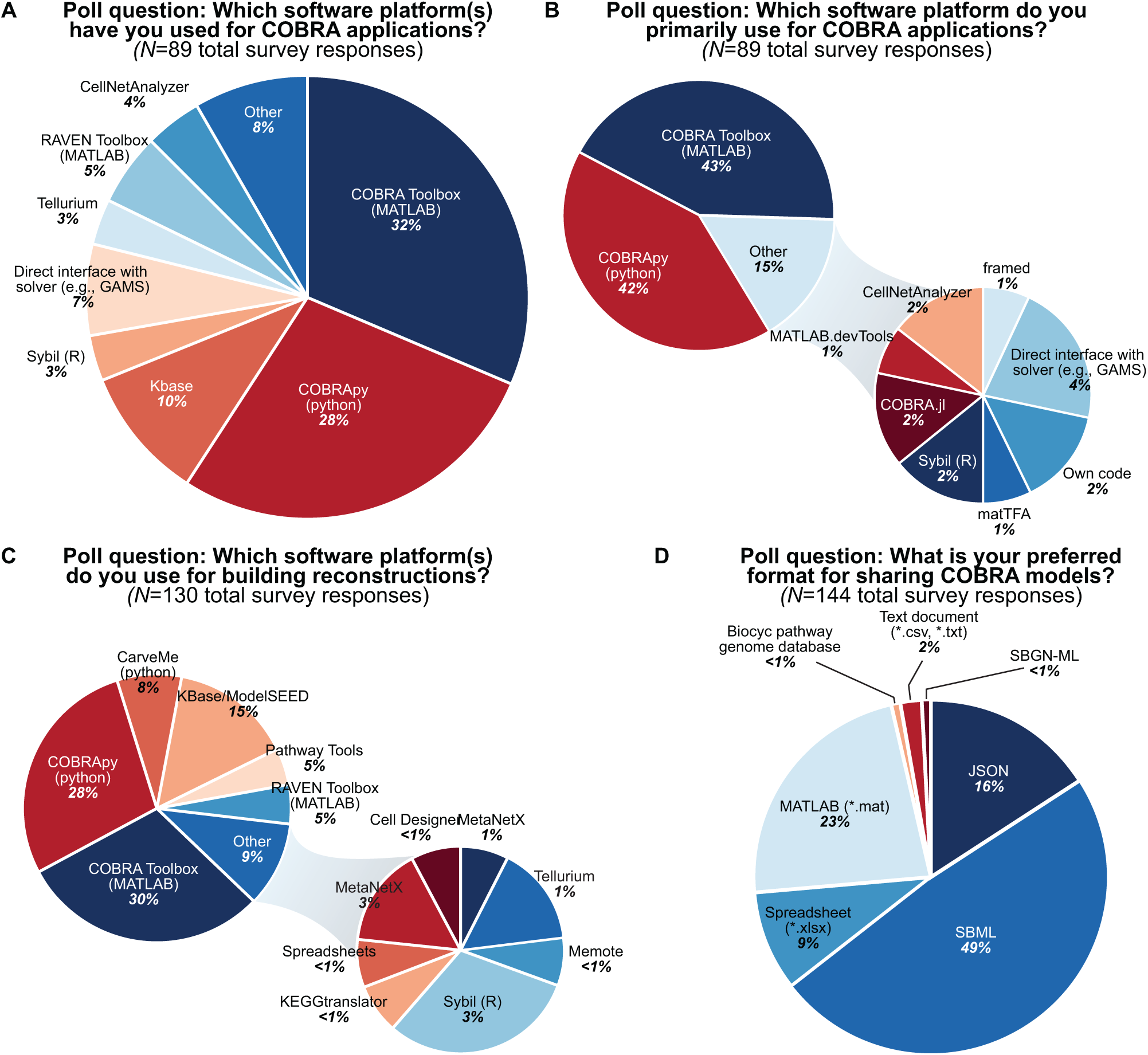
Poll results from the COBRA community survey. The survey was initially compiled and released at the 5th Annual Conference on Constraint-based Reconstruction and Analysis (COBRA, October 14-16, 2018); feedback from the conference was used to refine the survey, with an updated version later shared via social media (results are shown here; raw data provided in Data S1). The survey included 16 multiple choice and three open ended questions to summarize the field’s use and awareness of existing standards, as well as collect community-identified challenges. A total of 89 researchers completed the survey, representing different levels of expertise in the field; some questions permitted multiple responses (panels C and D). De-identified survey results can be accessed at: https://github.com/maureencarey/community_standards_supplemental.

**Table 1.** Resources for using community standards and software tools. Resources developed for broad applications in computational systems biology are denoted with an asterisk; unmarked resources are specific to the COBRA field.

| Resource | Description | Link/reference |
| --- | --- | --- |
| MIRIAM* | Minimum Information Required In the Annotation of bio-chemical Models | (Le Novère *et al.* 2005) |
| MIASE* | Minimum Information About a Simulation Experiment | (Waltemath *et al.* 2011) |
| Memote | MEtabolic MOdel TESTs (Lieven *et al.* 2018b) | https://memote.io |
| COBRA-related Google groups | Help for users of COBRApy, the python implementation of COBRA software (Ebrahim *et al.* 2013) | https://groups.google.com/forum/#!forum/cobra-pie |
|  | Help for users of the COBRA Toolbox, the MATLAB implementation of COBRA software (Heirendt *et al.* 2019) | https://groups.google.com/forum/#!forum/cobra-toolbox |
|  | Discussion forum of the systems modeling community of the International Society for Computational Biology (ISCB) | https://groups.google.com/forum/#!forum/sysmod |
| COBRA GitHub | Repository for COBRA software, includes issue and help pages | https://github.com/opencobra/ |
| COMBINE* | Community for coordinating standards for modeling in biology (umbrella organization for SBML, MIRIAM, MIASE, and more) | http://co.mbine.org |
| Kbase Help Board | Issue-tracking system to aid users to utilize tools and datasets | https://kbase.us/help-board/ |
| SBML Validator* | Tests the syntax and internal consistency of an SBML file | http://sbml.org/Facilities/Validator/ |
| SBML Test Suite* | Conformance testing system to test the degree and correctness of the SBML support provided in a software package | http://sbml.org/Software/SBML_Test_Suite |
| BiGG Models | Freely accessible database of GEMS (King *et al.* 2016) | http://bigg.ucsd.edu |
| BioModels* | Repository of mathematical models of biological and biomedical systems (Glont *et al.* 2018) | https://www.ebi.ac.uk/biomodels/ |
| MetaNetX | Platform for accessing, analyzing, and manipulating GEMS (Moretti *et al.* 2016) | https://www.metanetx.org |

Ultimately, SBML is just a serialization of a particular data model. The format of the serialization itself is not crucial; what matters is the format’s ability to represent the necessary data structures and whether information can be unambiguously encoded and made freely accessible. SBML enforces interpretable encodings. Further, these standards must be widely accepted to be easily used in multiple software tools. This pervasiveness is essential—especially for network reconstructions—where the same knowledgebase could prove useful in various applications, requiring multiple tools in a complex analysis pipeline.

### Model testing

There are different types of model evaluation processes. One important kind of evaluation is to ensure syntactical correctness (e.g., ensure a model is saved as a valid and machine-readable SBML with the SBML validator or test suite; see Table 1); however, syntactical correctness does not imply biological meaning or computational correctness. Thus, a model must first be checked for syntax, then evaluated for biological sense—a multi-step process. A recent effort to improve standardization in the COBRA community resulted in Memote, a set of MEtabolic MOdel TEsts (Lieven *et al*. 2018a) to increase reproducibility and model quality through model evaluation. Using either a command-line software or a web interface, users can generate a report to evaluate a reconstruction, including (1) namespace of components, (2) biochemical consistency, (3) network topology, and (4) versioning. Memote focuses on both the technical correctness (i.e., syntax) of a model while also providing metrics that can help users to evaluate the biological correctness of the model.

Namespaces are evaluated for metabolites, genes, and reactions to check annotations for coverage (e.g., how many metabolites have an InChI key?), consistency (e.g., how many metabolites have correct InChI keys?), and redundancy (e.g., how many metabolites also have additional identifiers?). Biochemical consistency is evaluated to verify the preservation of mass and charge across both individual reactions and the entire network. Tests evaluating network topology are used to evaluate the more subjective features of the model; these test results can be used to infer the quality of a reconstruction (i.e., manually curated versus an automated draft reconstruction) when combined with biological knowledge of the system. Memote also reports the state of the software and environment versions used by the reconstruction and during the process of testing the reconstruction. The recent development of such a community-defined testing suite should improve the rigor of the field, particularly by tailoring general systems biology resources to our specific use cases. We encourage the community to make the use of Memote an expected standard for newly published models.

### Model quality and content

Many of the standards used in the COBRA field were developed by interdisciplinary teams of modelers and software developers for broad use in the computational biological modeling field (Table 1); we can use these existing standards or adapt them for use in our field as done with Memote (Box 1). For example, minimum and recommended quality standards have been formulated and presented as a set of expectations for biological models and simulations through (1) the Minimum Information Required In the Annotation of biochemical Models (MIRIAM (Le Novère *et al*. 2005)) and (2) Minimum Information About a Simulation Experiment (MIASE (Waltemath *et al*. 2011)), respectively. However, engagement in the COBRA field in particular has been modest, likely due to community members’ lack of familiarity with these resources (Data S1) and the challenges associated with updating these recommendations with new data types and applications. In the following sections, we discuss potential challenges facing the widespread adoption of these standards in the COBRA field and possible solutions.

**Box 1** Pipeline for genome-scale metabolic network modeling, including existing and proposed standards. Reconstructions represent powerful tools for the interrogation and understanding of metabolism. Here we outline the modeling pipeline and the decisions made throughout this process, as well as existing and proposed standards.

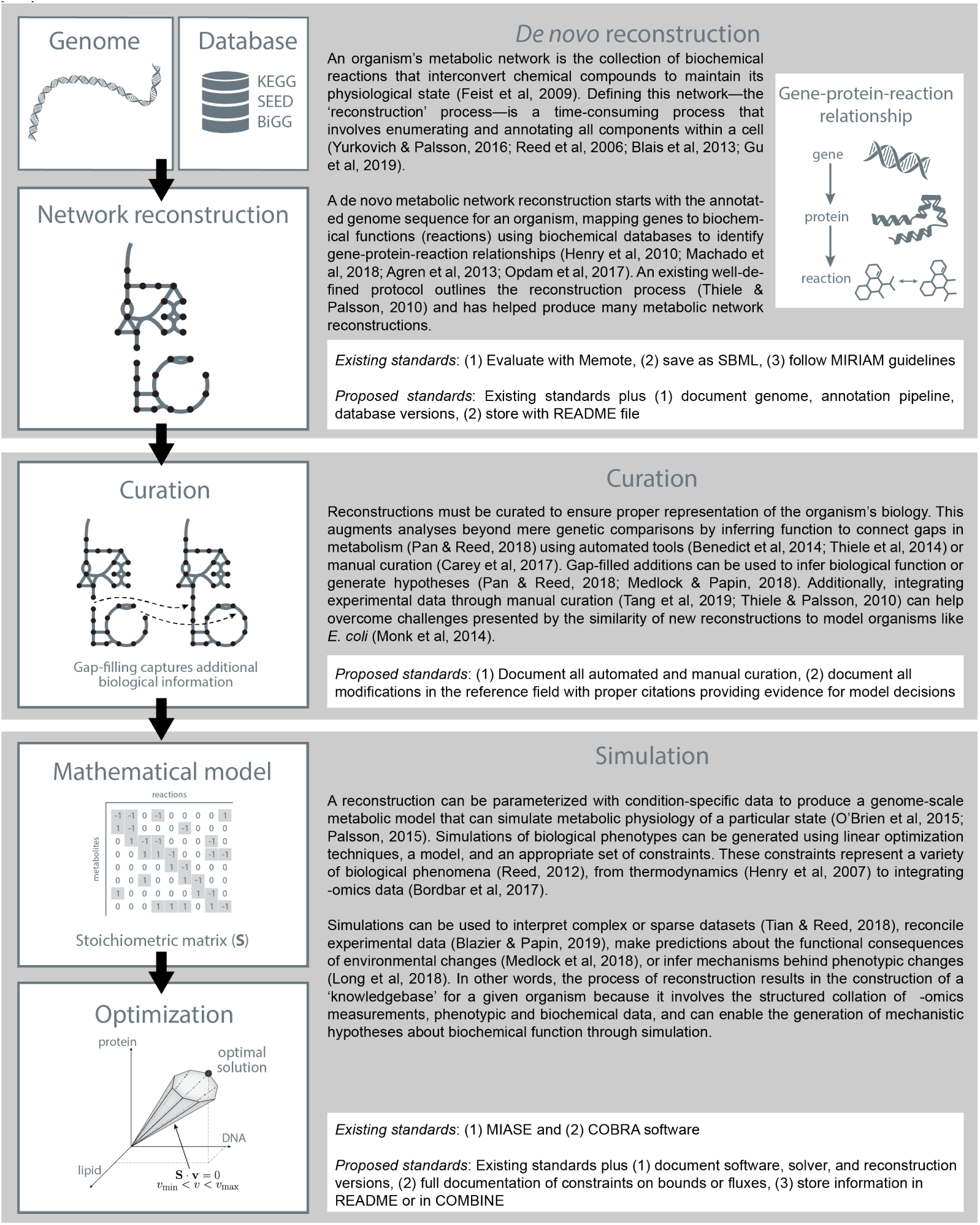

## CHALLENGES PREVENTING THE USE OF STANDARDS

Despite these efforts, many genome-scale metabolic network models fail to meet minimum standards and quality metrics. Ravikrishnan and Raman found that almost 60% of models had no standardized (i.e., interpretable) metabolite identifiers, 36% could not be evaluated for mass imbalances due to unstandardized formatting, and 35% did not contain gene-protein-reaction associations in the SBML file (Ravikrishnan and Raman 2015). This is a broad challenge throughout systems biology fields (Stanford *et al*. 2015). As a community, we must therefore ask why standards are not used more broadly if they enable the sharing, reuse, and evaluation of biological models and associated simulations. At the 5th Annual Conference on Constraint-based Reconstruction and Analysis (COBRA 2018), we surveyed the COBRA community regarding the use of community standards. This survey identified two major drivers for the lack of standardization in the COBRA field (full anonymized survey results provided in Data S1).

First, the responses identified several complex biologies that are not captured by current standards. For example, modelers of intracellular pathogen metabolism struggle to comply with nomenclature and mass balance when adding both pathogen and host biochemistry (Carey *et al*. 2017). Similarly, it is challenging to use the correct and sufficiently detailed nomenclature for biologically relevant tautomers and polymers (Data S1). While such issues will likely only be relevant in specific biological applications, it is vital for these edge cases to be addressable by community-adopted standards.

Second, users identified a set of novel analyses that current standards do not sufficiently support (Data S1). Current standards are inherently insufficient for novel techniques. Extensive community networks—such as modeling multiple members of the microbiota (Magnúsdóttir *et al*. 2017)—and modeling macromolecular expression mechanisms (Yang *et al*. 2018) represent current areas in metabolic modeling where some standards are currently lagging. Although standards must and do evolve as the field progresses, they inherently cannot capture the latest cutting-edge developments, just as biology textbooks cannot capture the latest scientific findings. This ‘lag’ in standardization is not field-specific (Hernandez *et al*. 2010), and such cutting-edge examples will likely only be identified in novel methods development. Both of these user-identified limitations require community-driven efforts to update standards as the field expands into new application areas and with novel analytic approaches.

We hypothesize that two additional factors play a role in these standardization challenges. First, biologists, modelers, and software developers are sometimes ‘siloed’ into separate communities and with distinct motivating factors (e.g., research interests, funding mechanisms). As a result, biologists and modelers are often not aware of relevant resources generated by software developers (Data S1). Our survey identified that fewer than 25% of researchers in the COBRA field were familiar with MIASE and only 56% were aware of MIRIAM (Data S1); these best practices cannot be used if they are not known. In turn, biological limitations—like those discussed above—might not be relayed to software developers focusing on a standard formulation. Thus, even community-driven efforts do not necessarily move laterally across subdisciplines. Second, as users, the lack of standardization often makes it easier to generate a novel reconstruction or analytic tool than to improve upon an existing version, further diversifying the set of existing approaches and amplifying the challenge of developing unifying standards.

## COMMUNITY-DRIVEN SOLUTIONS

One solution to increase the standardization in the field is to penalize noncompliant models through the manuscript review process; more than 85% of community-survey responders think this should occur (Data S1). However, there is little incentive against sharing a noncompliant reconstruction or even for failing to make a reconstruction publicly available. To remove these barriers, we suggest the field shifts to incentivize standardization, reuse, and markers of quality, and ultimately to improve communication amongst biologists, modelers, and software developers. Incentivizing standardization through funding models is inherently challenging: funding for science is evaluated in the short-term, whereas the benefits of software or resource development are observed on a longer time scale. We can integrate funding for infrastructure into applied projects to pair funding for the biological applications of genome-scale metabolic modeling and infrastructure development for the COBRA field.

Communication between those developing standards and those attempting to use standards is essential. One possible solution would be better mingling between the communities that design standards (e.g., SBML) and the communities that use them (e.g., COBRA), a solution that could be implemented through scientific meetings: each conference could have a dedicated keynote presentation by a representative from the other community, followed by a panel discussion led by the presenting representative. By maintaining clear contributing instructions for the COBRA software suites, associated analysis packages, and software for infrastructure, innovative solutions can be added to the standards by those who identify the cutting-edge fringe cases.

## RECOMMENDATIONS FOR STANDARDS

In response to some of the issues and challenges outlined above, we propose a set of guidelines to help improve the accessibility, content, and quality of metabolic network reconstructions—both for those creating reconstructions/models (Boxes 1 and 2) and those peer-reviewing reconstructions/models (Box 3). The suggestion of these standards was informed by panel discussions at the COBRA 2018 conference and from the community poll results (Data S1), as well as previous community efforts (Stanford *et al*. 2015). Our recommendations here represent field-specific implementation of the FAIR Data Principles (Wilkinson *et al*. 2016), a set of guidelines intended to improve reproducible research (Sansone *et al*. 2019).

First, focusing on the reconstruction process, we propose that a reconstruction metadata file is shared and includes model building information, such as the genome, database, and software versions (example README.md is provided in Data S2 or COMBINE archive in Additional file 2 of (Bergmann *et al*. 2014)). Although this information is likely in the original manuscript, this format would link the reconstruction to the reconstruction file.

Second, we encourage the use of version control and specific effort to document automated and manual curation. Version control can be implemented in multiple ways, mainly through a publicly available repository that includes all iterations or by making all versions publicly available and identifiable through clear naming conventions. Further, we propose that all curation efforts be documented in the reconstruction and explicitly include a literature reference and notes in the annotations field of a reaction.

Third, we emphasize the need for MIASE requirements (Waltemath *et al*. 2011) when sharing simulation results. These data about experimental data, constraints, and versioning can be stored in a COMBINE repository or the analytic code, if publicly available. Ultimately, a standardized format (like COMBINE) could enable minor advances in COBRA software to facilitate the re-implementation of a simulation.

## LOOKING AHEAD

Here, we have summarized existing standards in the COBRA field and identified challenges associated with both the development and compliance of software and model standards. We have proposed ‘checklists’ for use during both the reconstruction and peer review processes that will help improve the accessibility, content, and quality of metabolic network reconstructions. Additional community-inspired challenges and results from the COBRA community survey conducted in early 2019 are documented in the Data S1; we hope these examples will inspire new discussions and novel solutions.

There exist several open challenges for the field regarding the adoption of and development of new standards. We must embrace flexible standardization to facilitate their adoption and to build upon existing work. For example, although resources like MetaNetX (Moretti *et al*. 2016) and the BiGG Models database (King *et al*. 2016) facilitate the mapping of genes, reactions, and metabolites across the different namespaces, nomenclature discrepancies remain a challenge and sometimes result in redundant nonstandardized efforts. Another challenge is for community standard development to be derived from the community instead of in a top-down manner. While this organizational structure is currently in effect for the SBML community, it only functions if there is community participation—we need those who use the standards (i.e., modelers) to raise their hands and participate in the decision making process.

Ultimately, community adherence to standards will improve modeling reproducibility and better document the reconstruction process. We hope that the community embraces existing standards and our community-driven suggestions moving forward—both during the preparation of manuscripts and during the peer review process—and anticipate that compliance will increase the rigor of the field while simultaneously making it easier for scientists from other disciplines to build and use metabolic models.

**Box 2:** Proposed minimum standardized content for a metabolic network reconstruction. We propose that modelers use this list as a guide to help standardize accessibility, content, and quality; however, more comprehensive documentation and more interpretable and accessible information can only improve the usability and biological relevance of the shared reconstruction.

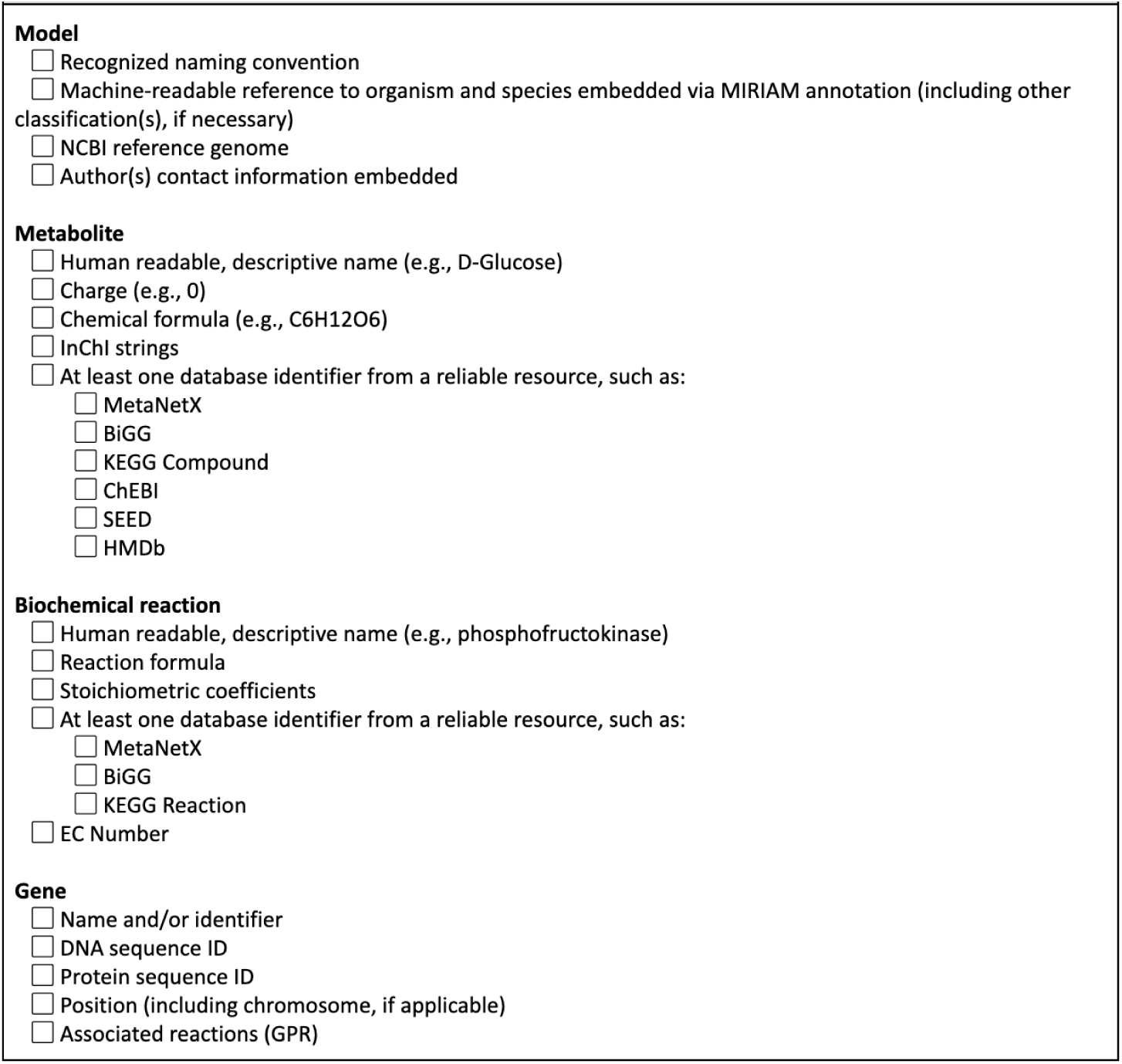

**Box 3:** Proposed checklist for reviewers. We propose that reviewers of manuscripts that include a novel metabolic network reconstruction use this list as a guide to help standardize accessibility, content, and quality.

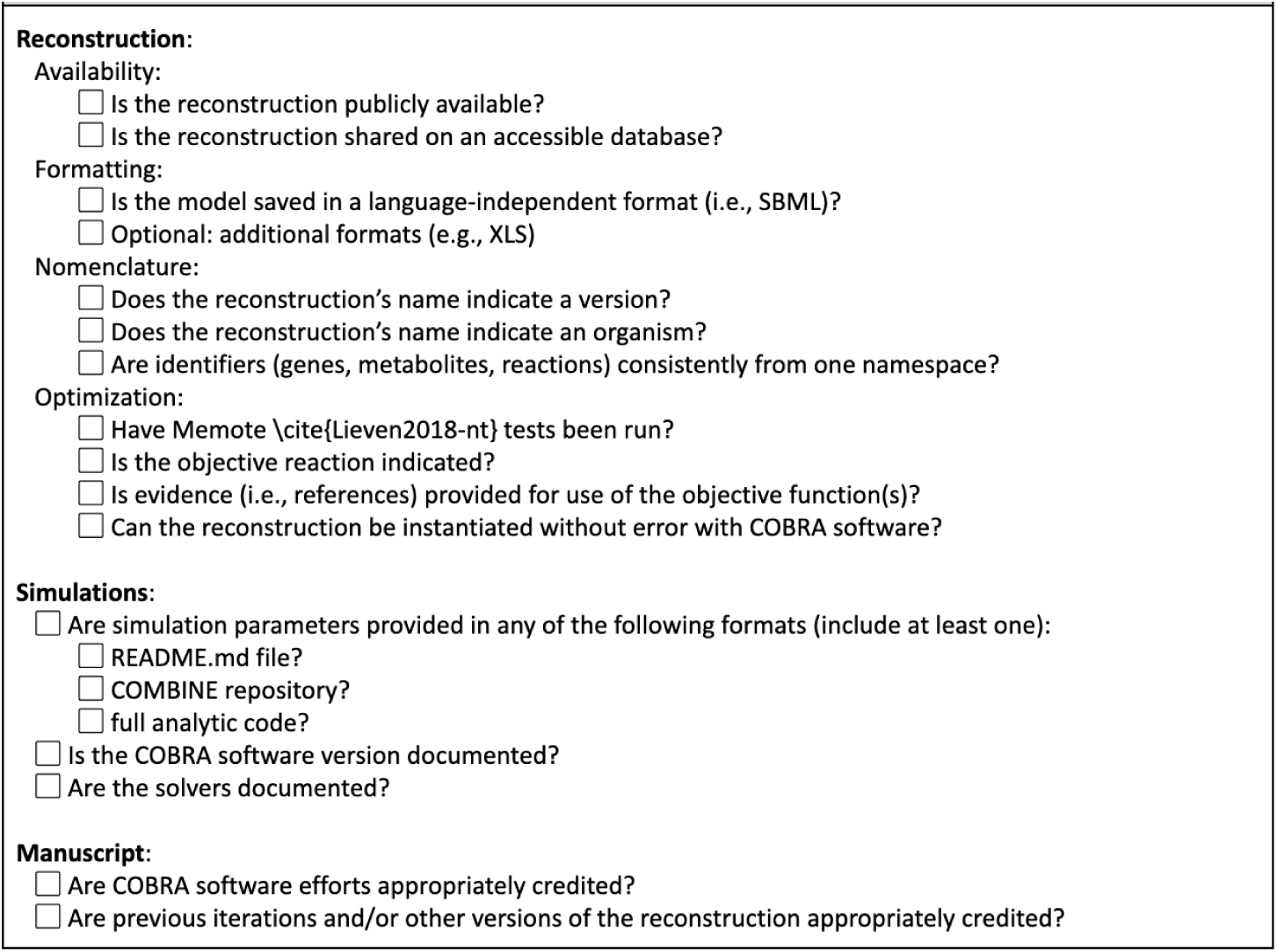

## ACKNOWLEDGEMENTS

The authors would like to thank those who contributed to the community survey. This work was funded by the Institute for Systems Biology’s Translational Research Fellows Program (J.T.Y.), the University of Virginia’s Engineering-in-Medicine Seed Funding (M.C. and J.P.), the PhRMA Foundation Postdoctoral Fellowship for Translational Medicine and Therapeutics (M.C.), and the National Institutes of Health (2R01GM070923-13 to A.D. and R01AT010253 to J.P.).

